# Adult Muscle Stem Cell Self-Renewal Induced by Endurance Exercise is Mediated by Inhibition of Mitochondrial Oxygen Consumption

**DOI:** 10.1101/845438

**Authors:** Phablo Abreu, Alicia J. Kowaltowski

## Abstract

Skeletal muscle stem cells (satellite cells) are well known to participate in regeneration and maintenance of the tissue over time. Studies have shown increases in the number of satellite cells after exercise, but their functional role in endurance training remains unexplored. Here, we found that injured muscles from endurance-exercised mice display improved regenerative capacity, demonstrated through higher densities of newly formed myofibers compared to controls, as well as lower inflammation and fibrosis. Enhanced myogenic function was accompanied by an increased fraction of satellite cells expressing self-renewal markers. Control satellite cells had morphologies suggestive of early differentiation, while endurance exercise enhanced myogenic colony formation. The beneficial effects of endurance exercise were associated with satellite cell metabolic reprogramming, including reduced mitochondrial respiration (O_2_ consumption) under resting conditions (absence of muscle injury) and increased stemness. During proliferation or activated states (three days after injury), O_2_ consumption was equal in control and exercised cells. Surprisingly, inhibition of mitochondrial O_2_ consumption was sufficient to enhance muscle stem cell self-renewal characteristics *in vitro*. Moreover, transplanted muscle satellite cells from exercised mice or cells with reduced mitochondrial respiration promoted a significant reduction in inflammation compared to controls. We propose that endurance exercise promotes self-renewal and inhibits differentiation in satellite cells, an effect promoted by metabolic reprogramming and respiratory inhibition, and which is associated with a more favorable muscular response to injury.

## Introduction

Satellite cells are the resident stem cells of the skeletal muscle, capable of regenerating the tissue throughout adulthood in response to insults (Yin et al., 2013). Regeneration occurs by activation of satellite cells, which can then differentiate and either fuse with existing multinuclear contractile myofibers and repair them, or generate new myofibers by fusing with other activated satellite cells.

Satellite cells express paired box protein 7 (Pax7), a critical regulator of muscle stem cell maintenance and activation. Indeed, loss of this protein leads to lack of satellite cell expansion and differentiation in both neonatal and adult muscles (von Maltzahn, 2013; Bazgir et al., 2017). Activation of these stem cells into committed myoblasts occurs due to extrinsic myogenic stimuli, and typically involves expression of the myogenic regulatory factor MyoD. The committed myogenic progenitors then actively proliferate to generate sufficient committed myoblasts, as well as produce differentiated myocytes, characterized by the expression of myogenin. Activated cells can also return to quiescence and self-renew, a process marked by Spry1 and Notch expression.

One of the most effective stimuli to induce satellite cell activation in adult muscles is exercise. Indeed, resistance exercise has long been known to activate and increase numbers of satellite cells, as well as to promote myogenesis (Darr and Schultz, 1987). Interestingly, however, the effects of aerobic endurance exercise on satellite cell populations and activation is less understood, and mixed results have been described in the literature (Mangan et al., 2014; Bazgir et al., 2017). This is an important point of study, since endurance exercise is well-known to improve skeletal muscle fitness (Jones and Carter, 2000), and is therefore expected to affect satellite cell activation, self-renewal and differentiation.

Mitochondrial function in differentiated myocytes is modified by endurance exercise, which promotes increased biogenesis of the organelle due to activation of the transcriptional co-regulator PGC-1, as well as upregulated mitochondrial quality control pathways (Baar et al., 2002; Balan et al., 2019.; Tang and Rando, 2014). Indeed, loss of mitochondrial mass and function occurs in aging, and may be a cause of age-related sarcopenia and frailty, which is partially prevented by endurance exercise (Rooyackers et al., 1996; Conley et al., 2000; Chabi et al., 2008, Balan et al., 2019).

Mitochondrial function is also determinant in stem cell function and differentiation (reviewed in Khacho and Slack, 2017). Although the type of mitochondrial functional alteration during commitment to differentiation is variable with stem cell fate (Forni et al., 2016), activation of oxidative phosphorylation is often required upstream of differentiation, while suppression is necessary for self-renewal in pluripotent stem cells (Khacho and Slack, 2017). Little is known, however, about how changes in mitochondrial bioenergetics are related to satellite cell activation, self-renewal or differentiation.

In this study, we investigated the effects of endurance exercise on skeletal muscle regeneration and satellite cell function. Surprisingly, we found that endurance exercise induces a significant repression of satellite cell oxidative phosphorylation, which stimulates self-renewal.

## Methods

### Animals

Two month old male C57BL/6 mice were purchased from Anilab, Brazil. All experimental procedures were conducted in agreement with Ethical Principles in Animal Research of the Brazilian College of Animal Experimentation (CONCEA). Protocols were reviewed and approved by the local Animal Care and Ethics Committee (protocol number 137/2017). Mice were housed 3-4 per cage and maintained on a 12:12 h light-dark cycle in a temperature-controlled environment (22°C) with *ad libitum* access to standard laboratory chow (Nuvital Nutrients, Curitiba, Brazil) and water.

### Incremental speed test and endurance exercise protocol

Mice were submitted to incremental speed exercise testing on a motor treadmill before and after the experimental period of endurance training. After adaptation to the treadmill environment for 1 week (10 min per session), mice were placed on the treadmill at 0% inclination and allowed to acclimatize for at least 10 min, then incremental speed increases started at 10 meters · min^−1^ during 10 min and were increased by 3 meters · min^−1^ every minute. Mice that could no longer run for over 1 min were considered exhausted and removed from the treadmill. This incremental speed test provided the maximal running duration (minutes), distance (meters) and velocity (meters · min^−1^). Peak workload was measured at the end of the test.

Endurance exercise was randomly assigned for sedentary (control) and exercised mice as 60% of the maximum intensity (maximal aerobic capacity) reached during the incremental speed test, as previously described (Ferreira et al. 2007; Abreu et al. 2012 and 2016; Campos et al. 2017). Exercised mice performed moderate-intensity running exercises on a motorized treadmill for 5 weeks (from the 8^th^ to 13^th^ weeks of age), 5 days a week, 60 min per day. The running velocity during the period of physical training was gradually increased to 60% of maximal aerobic intensity (described above). During 5 weeks, treadmill-running skills were maintained in sedentary (control) mice for 5 min (10 meters · min^−1^), twice a week, to avoid any interference of treadmill stress. This activity in control animals did not alter maximal exercise capacity (Abreu et al., 2016). Body weights and food consumption were recorded weekly.

Forty-eight hours after the last exercise at 5 weeks of endurance training, run capacity (maximal running times, velocities and distances) was evaluated in all mice (Medeiros et al. 2004; Ferreira et al. 2007; Abreu et al., 2012 and 2016; Campos et al. 2017). After this session, the animals were submitted to live animal experiments as described below or anesthetized with isoflurane and sacrificed by cervical dislocation for tissue or cell collection. Tissues were quickly removed, weighed, immediately frozen in liquid nitrogen, and stored at −80°C.

### Indirect calorimetry, spontaneous physical activity and maximal oxygen consumption (VO_2_max) determination

A room with restricted access was dedicated for this experiment. Oxymax, (Columbus Instruments) and TSE Systems were used to monitor real time changes in gas concentrations. Animals were places in individual 30 cm diameter cages (n = 4, control and n = 4, exercised). Each mouse was acclimatized to an individual chamber 24 h before the measurements, and data were collected every 20 min for 24 h. Results shown were collected during the active period, corresponding to the early dark hours.

Infrared detection was used to measure locomotor activity in parallel with the indirect calorimetry assessments. Data were recorded for at least one full circadian period (24 h), and results shown were collected during the early dark hours. Movements were added for every minute and detected simultaneously on all three axes: forward and backward (x), side to side (y), and up and down (z).

Coupling of the treadmill with adjustable intensity to indirect calorimetry and an open-circuit calorimeter allowed for the assessment of maximal oxygen consumption (VO_2_max). Prior to the experiment, a familiarization session was run. Gases were measured every 10 sec. Average VO_2_ and running distance were calculated for each mouse (n = 10, control and n = 12, exercised).

### Muscle lesions

For muscle regeneration studies, injury was induced 24 hours after the last endurance exercise session. Mice (n = 4, control and n = 4, exercised) were anesthetized by inhalation using isoflurane. The belly region of the tibialis anterior muscle was injected intramuscularly with 50 μL 1.2% w/v barium chloride. Seven days after the injury, the animals were anesthetized with isoflurane and sacrificed by cervical dislocation. The muscles were quickly removed and stained.

For satellite cell activation studies, muscle injury was induced 24 hours after the last exercise session. Mice were anesthetized, and the belly region of the tibialis anterior, gastrocnemius, quadriceps and triceps brachii muscles were injected intramuscularly with 10 μL 1.2% w/v barium chloride. Three days after the injury, the animals were sacrificed, skeletal muscles were quickly removed, and satellite cells were isolated as described below.

### Stains and microscopy

For hematoxylin and eosin (H&E) staining analysis, injured muscles were embedded into tragacanth and Tissue-TEK® OCT solution and frozen in liquid nitrogen. Sections (10 μm thick) were obtained from the mid-belly region of the injured muscle (n = 4, control and n = 4, exercised) using a Leica CM3050 cryostat (Wetzlar, Germany). Sections were stained using the H&E method (Wittekind, 2003) and photographed using a Nikon Eclipse E1000 microscope coupled to a DXM 1200 camera. Five non-overlapping images from the central region of each section were analyzed using the cell counter plugin for Image J Software. Mean cross-sectional areas were determined by tracing 25 adjacent fibers, with a total of 500 fibers per group. Analysis was performed blinded.

Picrosirius Red was used to evaluate collagen content in injured muscles, as described by Junqueira et al. 1979. Sections were photographed using a polarized light microscope and images were captured on a JVC TK-1280E digital color video camera coupled to the microscope. A section of muscle fibers surrounded by perimysium, containing 10-100 fibers (n = 4, control and n = 4, exercised), was imaged as previously reported (Junqueira et al. 1979). Analysis was performed blinded.

### Dual x-ray absorptiometry measurements

Forty eight hours after the last endurance exercise session, mice (n = 4, control and n = 4, exercised) were scanned using the FX Pro *in vivo* imaging system (Akron, Ohio, USA). A combination of xylazine (10 mg/kg) and ketamine (90 mg/kg) was used for sedation. Fat area was measured in the abdomen (top of the pelvis to the lowermost rib) and calculated as abdominal fat divided by the total abdomen tissue. Lean area was evaluated in the upper hindlimbs.

### Muscle stem cell isolation, culture and clonal analysis

Muscle stem cells were isolated from intact extensor digitorum longus, gastrocnemius, quadriceps, soleus, tibialis anterior and triceps brachii muscles, as previously described (Cerletti et al, 2008; Conboy et al, 2003; Sherwood et al, 2004). Briefly, muscles were removed, minced, mechanically dissociated and filtered twice with 70 mm strainers. The resulting tissue was centrifuged and digested with 2% type II collagenase, 0.25% trypsin and 0.1% DNAase. Primary myoblasts were purified by 2-3 pre-plating steps (Gharaibeh et al. 2008) and cultured in Dulbecco’s Modified Eagle Medium (DMEM) containing 1% penicillin and 20% fetal bovine serum. In purity checks, 84.6% of the isolated cells with DAPI-stained nuclei were Pax7 positive, as assessed by immunofluorescence (described below).

For clonal analysis, cells were grown for 168 h on Matrigel-coated 6-well plates (5.000 cells per well), and then fixed with with 3.8% paraformaldehyde, dried for 24 h, died with 1% crystal violet and washed. After 24 h, clone colonies (clusters of approximately 50 cells) were manually counted.

### Immunofluorescence

Satellite cells from control and exercised mice were plated for 96 h on Matrigel-coated 12-well plates (100.000 cells per well), then fixed with 3.8% paraformaldehyde for 10 min, permeabilized with 0.3% Triton X-100 and blocked in 1% BSA, 0.3% Triton X-100, and 0.1% sodium azide in PBS for 2 h at room temperature. Cells were incubated with primary antibody (Pax7 monoclonal mouse IgG1, Abcam 199010; 1:50) at 4°C overnight followed by 4,6-diamidino-2-phenylindole (DAPI) for 1 h and fluorescent-labeled secondary antibodies (goat anti-mouse IgG conjugated to Alexa Fluor® 488; 1:1000). Images were captured with Coolsnap HQ CCD camera (Photometrics) driven by IP Lab software (Scanalytics Inc) using a Leica DMI 6000B fluorescent microscope (Mannheim, Germany). Quantification used the cell counter plugin for Image J software.

### Morphological analysis

Satellite cells from control (n = 8) and exercised (n = 8) mice were cultured for 96 h on Matrigel-coated 12 well plates (100.000 cells per well). Images were acquired on a Nikon Eclipse TS100 microscope and quantification of spindle-like and elongated morphology was conducted using the cell counter plugin for Image J.

### Metabolic flux analysis

Oxygen consumption rates (OCR) and extracellular acidification rates (ECAR) of muscle stem cells were measured using an XF24 Bio analyzer (Seahorse Bioscience), as described previously (Forni et al. 2016, Cerqueira et al., 2016). Cells were seeded in Matrigel-coated XF24 24-well micro plates at 20.000 cells/well for 120 h. Before the assay, the media was removed and replaced by 500 mL of low glucose (5 mM) assay medium, without sodium bicarbonate, to allow for ECAR measurements. ATP synthesis-linked O_2_ consumption and proton leak-driven respiration were determined by the addition of oligomycin (1 μg/mL). The uncoupler 2,4-dinitrophenol (2,4-DNP, 200 μM) was added to promote maximal respiratory capacity. Rotenone (1 μM) and 1 μg/mL antimycin A were added to ablate mitochondrial O_2_ consumption. All respiratory modulators were used at concentrations determined through preliminary titration analyses (Forni et al. 2016, Cerqueira et al., 2016).

### RNA extraction and real-time polymerase chain reaction (PCR) analysis

Cells were cultured on 6-well plates (250.000 cells per well) for 120 h. For mtDNA quantifications, DNA samples were prepared using a DNeasy® Blood & Tissue Kit (Qiagen). Real time quantitative PCR (qPCR) was performed using SYBR Green PCR Master Mix (Applied Biosystems). The following genes were analyzed: Hypoxanthine Phosphoribosyltransferase (HPRT) for nDNA and Mitochondrially Encoded NADH:Ubiquinone Oxidoreductase Core Subunit 1 (MT-ND1) for mtDNA copy number. Fold changes were calculated by the 2−ΔΔCT method. The primer sequences are presented in Supplementary Table 1.

Total RNA was purified using Trizol reagent (Invitrogen Life Technologies, Rockville, MD, USA), and quality-checked using 260/230 nm and 260/280 nm scores. Equivalent contents of RNA were reverse-transcribed using a Super-Script III cDNA Synthesis Kit (Invitrogen). The cDNA synthesized was stored at −20°C prior to the real-time PCR assay. Amplification was performed using Platinum® SYBR® Green qPCR SuperMix UDG (Invitrogen Life Technologies, California, USA) and evaluated by real-time PCR using the Rotor Gene 3000 apparatus (Corbett Research, Mortlake, Australia). Primer sequences were designed using information contained in the Gene Bank of the National Center for Biotechnology Information (NCBI). Gene expression was quantified using qBase software, as described previously (Hellemans et al. 2007). mRNA expression was normalized using the internal control gene HPRT (Nicot et al. 2005). Fold changes were calculated by the 2−ΔΔCT method. Primers were selected by Primer ban (https://www.exxtend.com.br); sequences are presented in Supplementary Table 1.

### Muscle stem cell transplantation

The tibialis anterior muscles of the hosts, 14-week-old male WT mice, were prepared by BaCl_2_ injury as described above, one day before transplantation; then 20.000 freshly isolated muscle stem cells from control or exercised mice in 30 μL media were slowly injected into the mid-belly of the muscle of a recipient anesthetized host (control) mouse. Seven days after transplantation, the muscles were quickly removed for microscopy.

### Statistical analysis

Results are presented as means ± SEM and were analyzed using GraphPad Prism Software 4.0. Data were compared using t-tests. Asterisks denote statistically significant differences versus control. Grubb’s test was used to exclude outliers.

## Results

### Endurance exercise increases performance and promotes a shift in substrate use

Mice were submitted to an endurance exercise protocol for 5 weeks (Fig. 1A), a time span sufficient to observe many of metabolic phenotypes associated with exercise (Ferreira et al. 2007; Abreu et al. 2012; Abreu et al., 2016; Campos et al. 2017). Their performance was then evaluated 48 h after the last exercise training session. Consistent with prior reports, exercised mice presented an increase in running time (Fig. 1B), speed (Fig. 1C) and distance (Fig. 1D) in the run-to-exhaustion test. Exercised mice also displayed a shift in energy substrate usage at rest analyzed through indirect calorimetry, as evidenced by increased O_2_ consumption (Fig. 1E) and eat production (Fig. 1G). Interestingly, spontaneous physical activity (Fig. 1F) was also increased in exercised mice (Speakman, 2013). Performance was also measured under run-to-exhaustion conditions. Oxygen consumption (Fig. 1H) and was again higher in exercised animals. Blood glucose monitoring during the run-to-exhaustion test showed more steady blood glucose levels in exercised animals (Fig. 1I; Abreu et al. 2016; Fan et al. 2017). Overall, these data indicate that the exercise regimen promoted enhanced performance associated with metabolic modifications.

**Fig 1.**
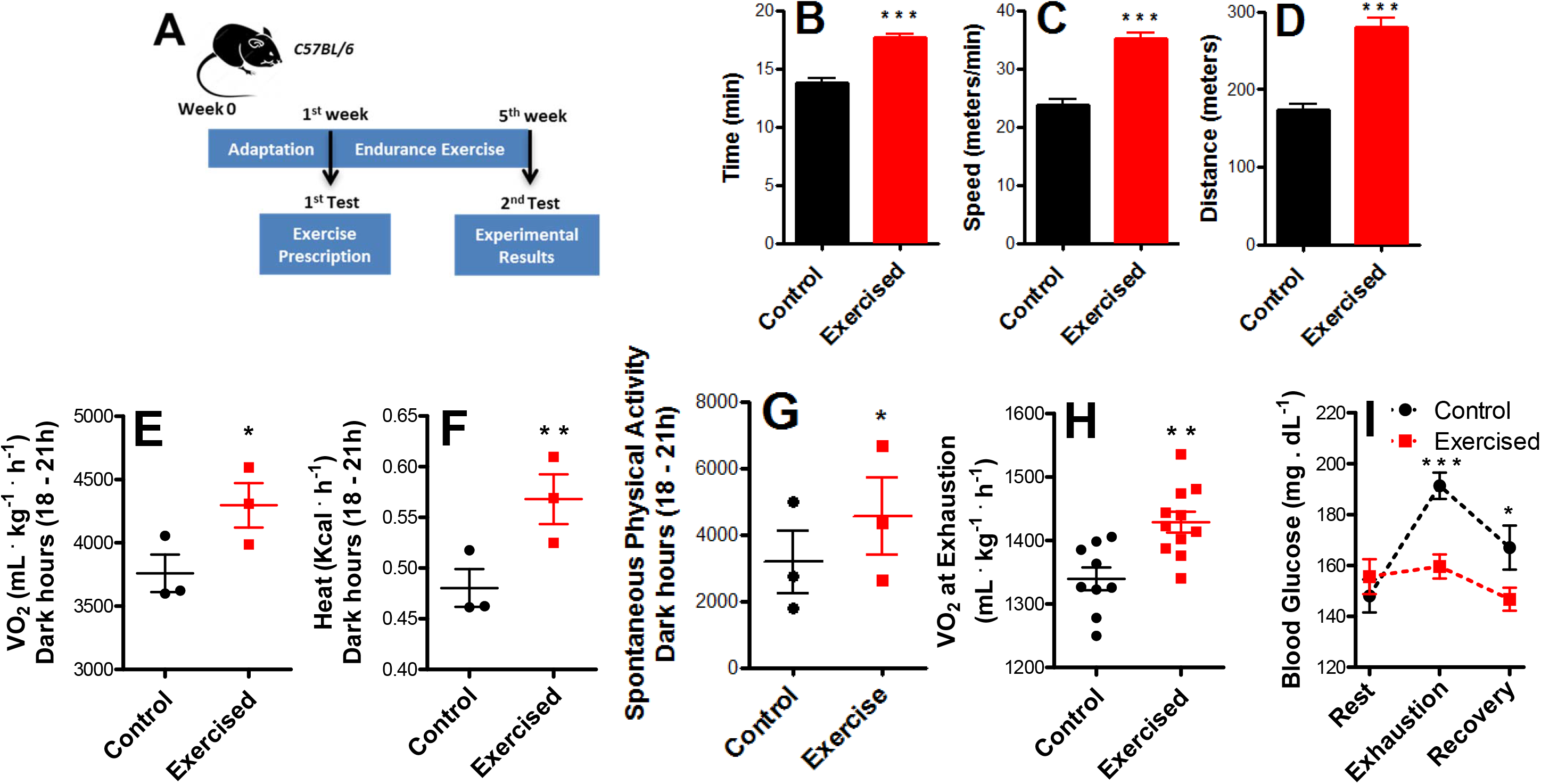
Endurance exercise induces a performance-dependent shift and changes energy substrate usage. A: Schematic panel. B: Running time. C: Running speed. D: Distance covered in the run-to-exhaustion test. E: O_2_ consumption during the dark hours (18-21 h) measured by indirect calorimetry. F: Heat production during the dark hours measured by indirect calorimetry. G: Spontaneous physical activity during the dark hours. H: O_2_ consumption at exhaustion. I: Resting blood glucose, blood glucose during run-to-exhaustion, blood glucose at 10 min rest after exhaustion. Data represent averages ± SEM and were compared using t-tests. Asterisks denote statistically significant differences (*p<0.05, **p<0.01, ***p<0.001) versus control.

Endurance exercise resulted in body mass reduction (Fig. 2A) that was not the sole result of changes in food consumption (Fig. 2B). Dual X-ray absorptiometry scans were used to measure body composition (Fig. 2C, representative images) and revealed a significant decrease in the relative area occupied by adipose tissue in this exercised group (Fig. 2D), with no difference in lean body composition (Fig. 2E). Indeed, muscle weights (Fig. 2F) did not change, nor were there any modifications in cross sectional areas of the soleus muscle (in which oxidative fibers are predominant) or tibialis anterior muscle (mostly glycolytic; Figs. 2G-I). Other organs such as the heart and liver also did not present altered mass (results not shown), suggesting weight loss is specific to fat depots.

**Fig 2.**
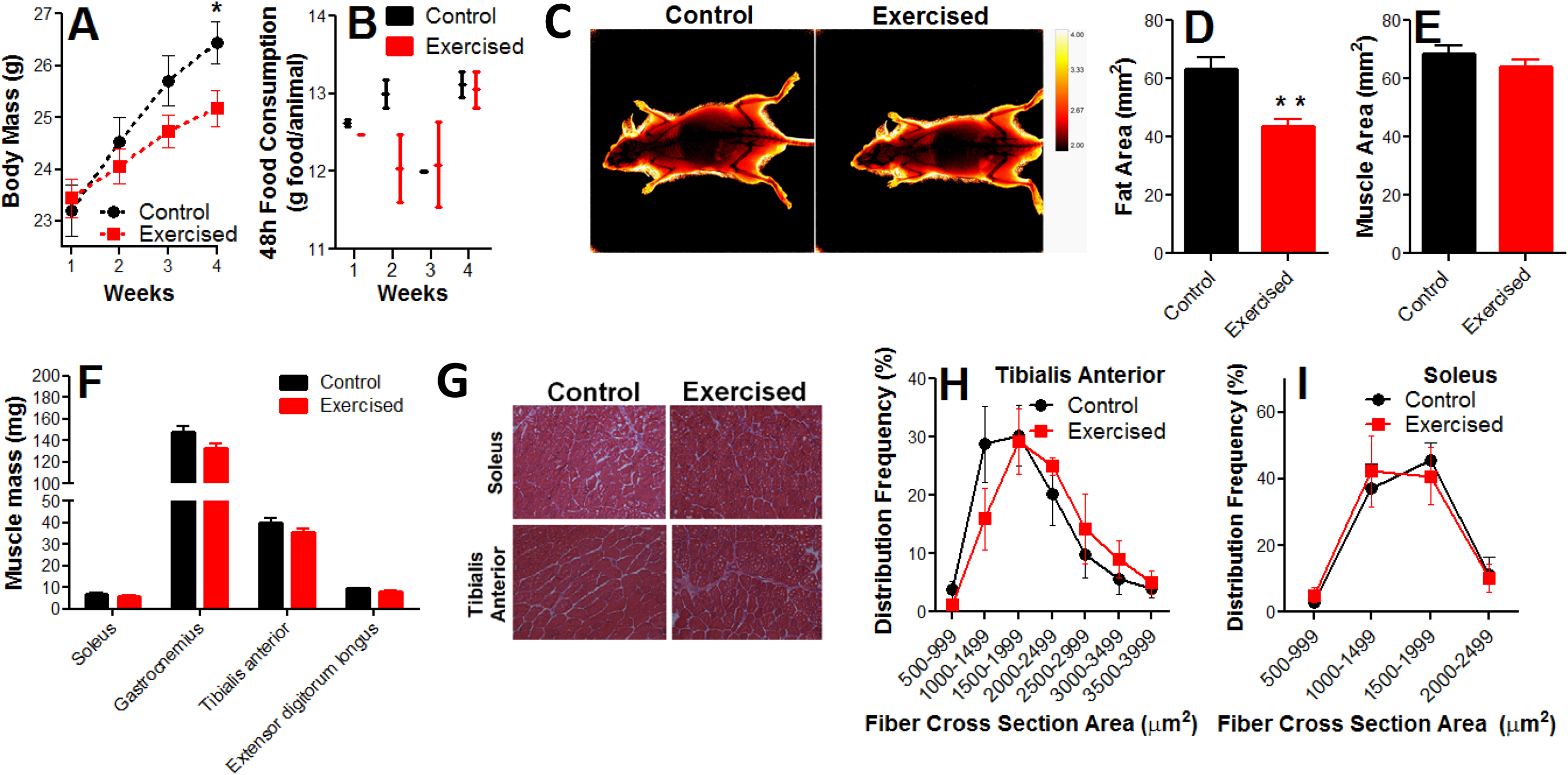
Body fat content is decreased by endurance exercise, while muscle content is preserved. A: Body mass changes. B: Food consumption. C: Dual X-ray absorptiometry scans, representative image. D: Fat area. E: Muscle area. F: Muscle mass. G: Representative images of cross sectional areas. H and I: Frequency distribution of the cross sectional areas of the fibers. Data represent averages ± SEM and were compared using t-tests. Asterisks denote statistically significant differences (*p<0.05, **p<0.01) versus control.

### Endurance exercise enhances skeletal muscle repair

Endurance exercise has been shown to protect against loss of satellite cells during aging, as well as increase their number and capacity to differentiate (Kurosaka et al. 2009; Shefer et al. 2010; Cisterna et al. 2016). We thus verified if satellite cell-mediated muscle regeneration after injury (Fig. 3A) was affected by endurance exercise. H&E microscopies of uninjured and injured tibialis anterior (TA) muscles from control (sedentary) or exercised animals are shown in Fig. 3B, and uncover no changes in cross-section area measurements (Fig. 3C). However, injured muscles from exercised mice had improved regenerative capacity, as demonstrated by the higher density of newly formed myofibers, which present central nuclei (Fig. 3D). Exercise also decreased the extent of inflammatory cell infiltration (Fig. 3E). Indeed, Picrosirius red staining (Fig. 3F), used to quantify the fibrous tissue area (Fig. 3G), demonstrates that exercised animals showed lower collagen content. We also quantified neutral lipid accumulation (results not shown), which was unchanged. Overall, these results show that the exercised muscle is preconditioned against injuries, resulting in higher fiber formation, lower inflammation as well as prevention of fibrosis.

**Figure 3.**
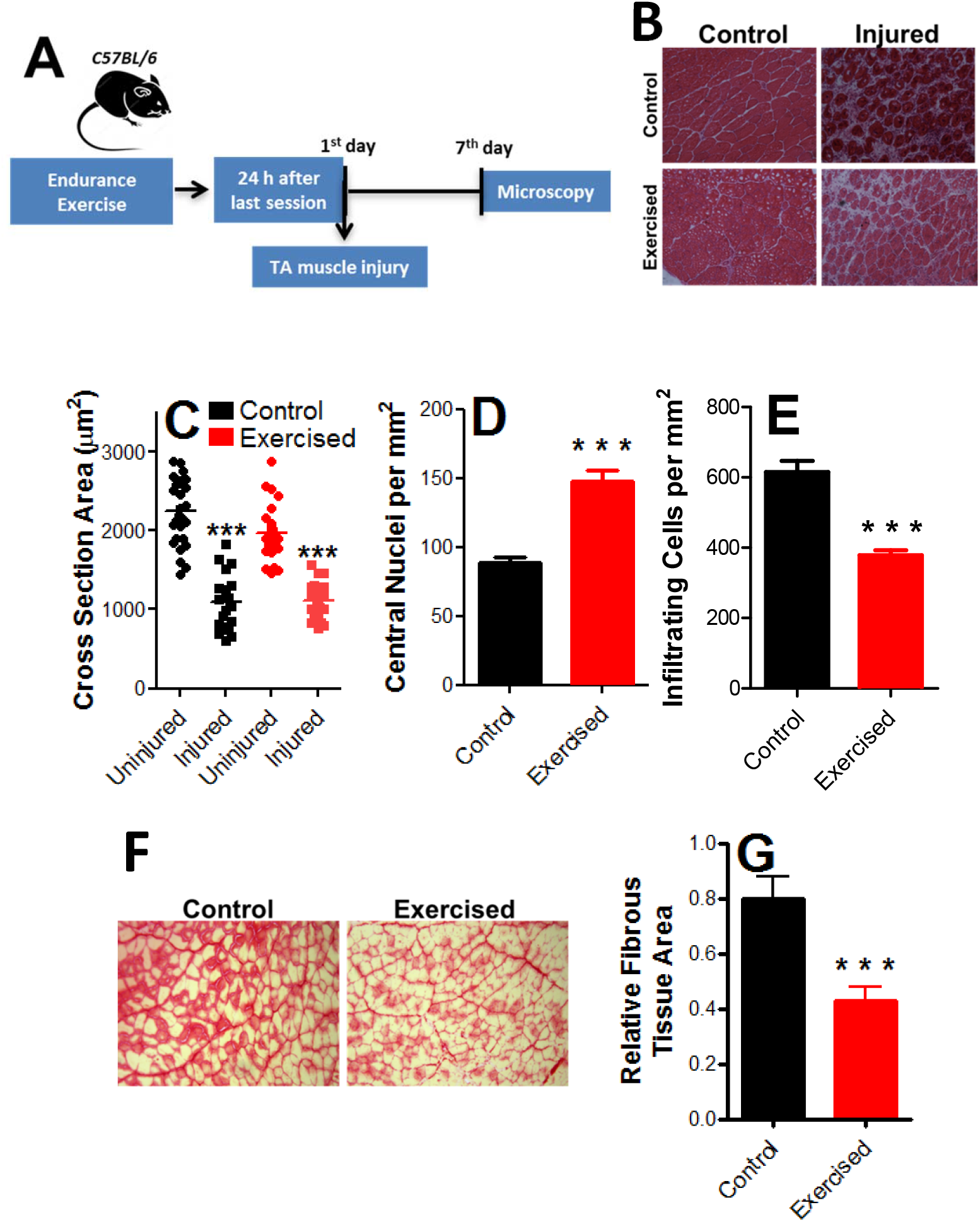
Aerobic fitness enhances skeletal muscle repair and promotes anti-inflammatory and anti-fibrotic effects. A: Schematic panel. B: Illustrative images of fiber cross-sections. C: Cross-sectional areas of control and exercised samples with and without injury. D: Centrally nucleated (newly-formed) myofiber counts in injured muscles. E: Infiltrating inflammatory cell counts in injured muscles. G: Picrosirius red-stained injured muscles, representative images. H: Relative fibrous area quantification. Data represent averages ± SEM and were compared using t-tests. Asterisks denote statistically significant differences (***p<0.001) versus controls.

### Endurance exercise enhances satellite cell self-renewal

The higher density of newly formed fibers and lower inflammation in endurance-exercised mice suggest changes in satellite cell population (Shefer et al. 2010; Cisterna et al. 2016). We thus isolated and plated satellite cells from exercised and control animals (Fig. 4A). Cell preparations presented high purity, as indicated by immunofluorescence assays for the satellite cell marker Pax7 (see Methods, Sacco et al., 2008). After 120 h of growth in culture, satellite cells were proliferative and at 70-80% confluence. At this time point, the expression of Pax7, MyoD1 (a marker for satellite cell activation) and CXCR4 (a satellite cell membrane marker) were significantly enhanced in cells isolated from exercised animals (Fig. 4B). Self-renewal markers Sirt1 (which regulates metabolic state and proliferation, Ryall et al., 2015, Myers et al., 2019), and HIF1α (Yang et al., 2017) were also increased by exercise, as were Spry1, Notch1 and Hey1, which signal myoblast return to quiescence, maintaining the satellite cell pool and long-term muscle integrity (Ryall et al., 2015). FOXO3a expression (a quiescence marker) was not modified (Fig. 4B). Overall, this expression profile suggests endurance exercise promotes activation, but prevents final satellite cell differentiation, maintaining myoblast self-renewal or return to quiescence.

**Fig 4.**
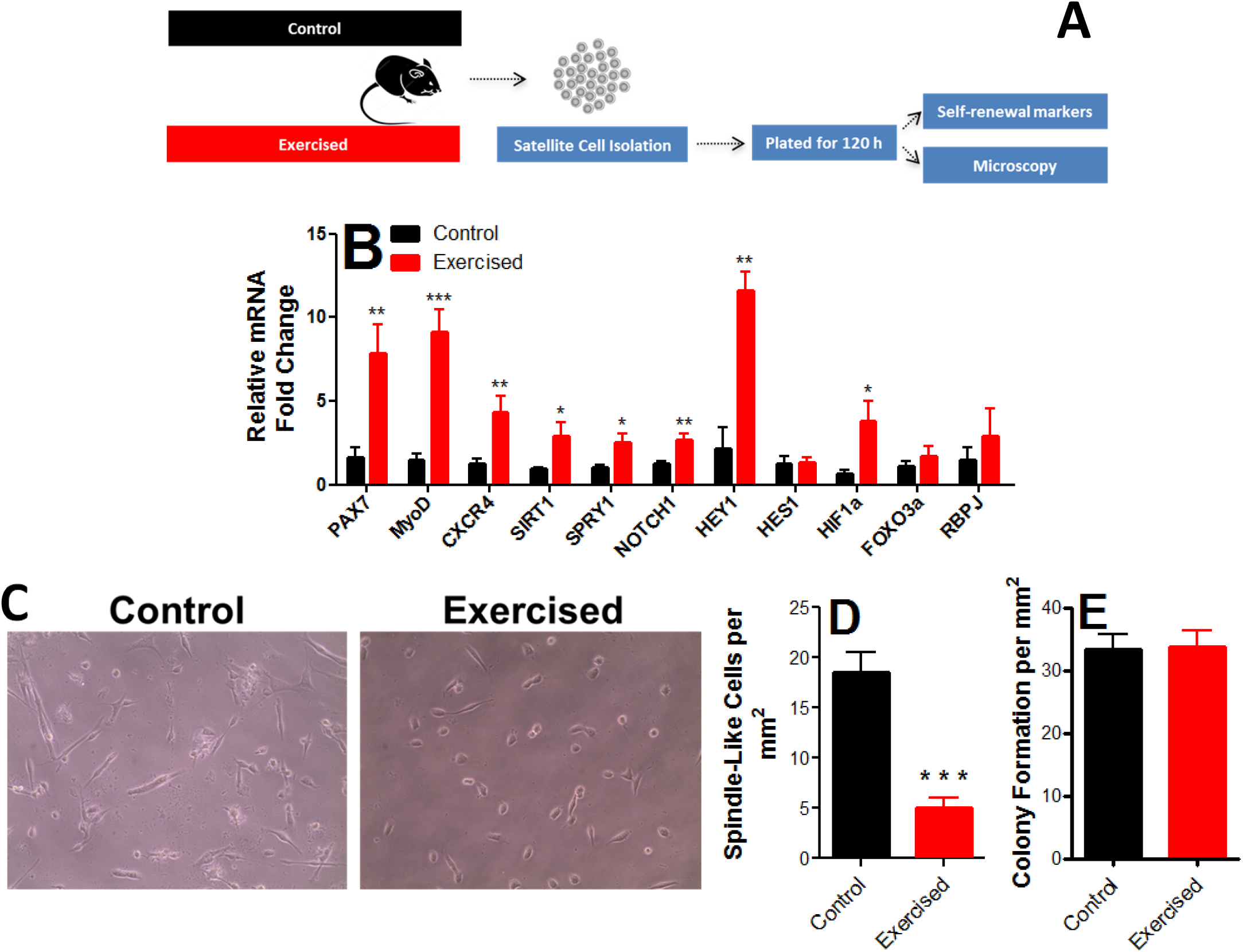
Endurance exercise enhances self-renewal and quiescence markers in satellite cells. A: Schematic panel. B: Expression of Pax7 (a satellite cell-specific marker which increases upon activation), MyoD1 (an activation marker), CXCR4 (a satellite cell marker), Sirt1 (a self-renewal marker), Spry1 (a return-to-quiescence marker), Notch1, Hey1, Hes1 and HIF1α (self-renewal markers) FOXO3a (quiescence marker) and Rbpj (involved in Notch1 signaling). C: Representative images of satellite morphology. D: Quantification of elongated, spindle-like (more differentiated), cells. E: Myogenic colony formation. Data represent averages ± SEM and were compared using t tests. Asterisks denote statistically significant differences (*p<0.05, **p<0.01, ***p<0.001) versus control.

To functionally assess the differentiation of these cells, cell morphology was evaluated (Fig. 4C-D). Control satellite cells from sedentary animals showed spindle-like, elongated, morphology, suggestive of early differentiation and loss of stemness, while endurance exercise promoted a more undifferentiated myoblast morphology. Despite changes in differentiation markers and morphology, no difference in colony formation ability was observed between exercised and control cells (Fig. 4E). Overall, our results suggest that endurance exercise promotes satellite cell self-renewal and quiescence, but inhibits differentiation, without affecting cell proliferation.

### Endurance exercise reduces satellite cell O_2_ consumption rates under physiological, but not activated states

There is literature evidence that satellite cell cycle regulation involves changes in energy metabolism, with modulation of substrate use (Zhang et al. 2016; Cerletti et al. 2012; Ryall et al. 2015; Rodgers et al. 2014). However, mitochondrial oxidative phosphorylation, which is a key regulatory factor in stem cell differentiation (Khacho and Slack, 2017; Forni et al., 2016), has not been directly assessed in the context of satellite cell modulation. We used a Seahorse Extracellular Flux analyzing system to assess mitochondrial bioenergetics in intact satellite cells isolated from control or endurance exercised animals (Fig. 5A).

**Fig 5.**
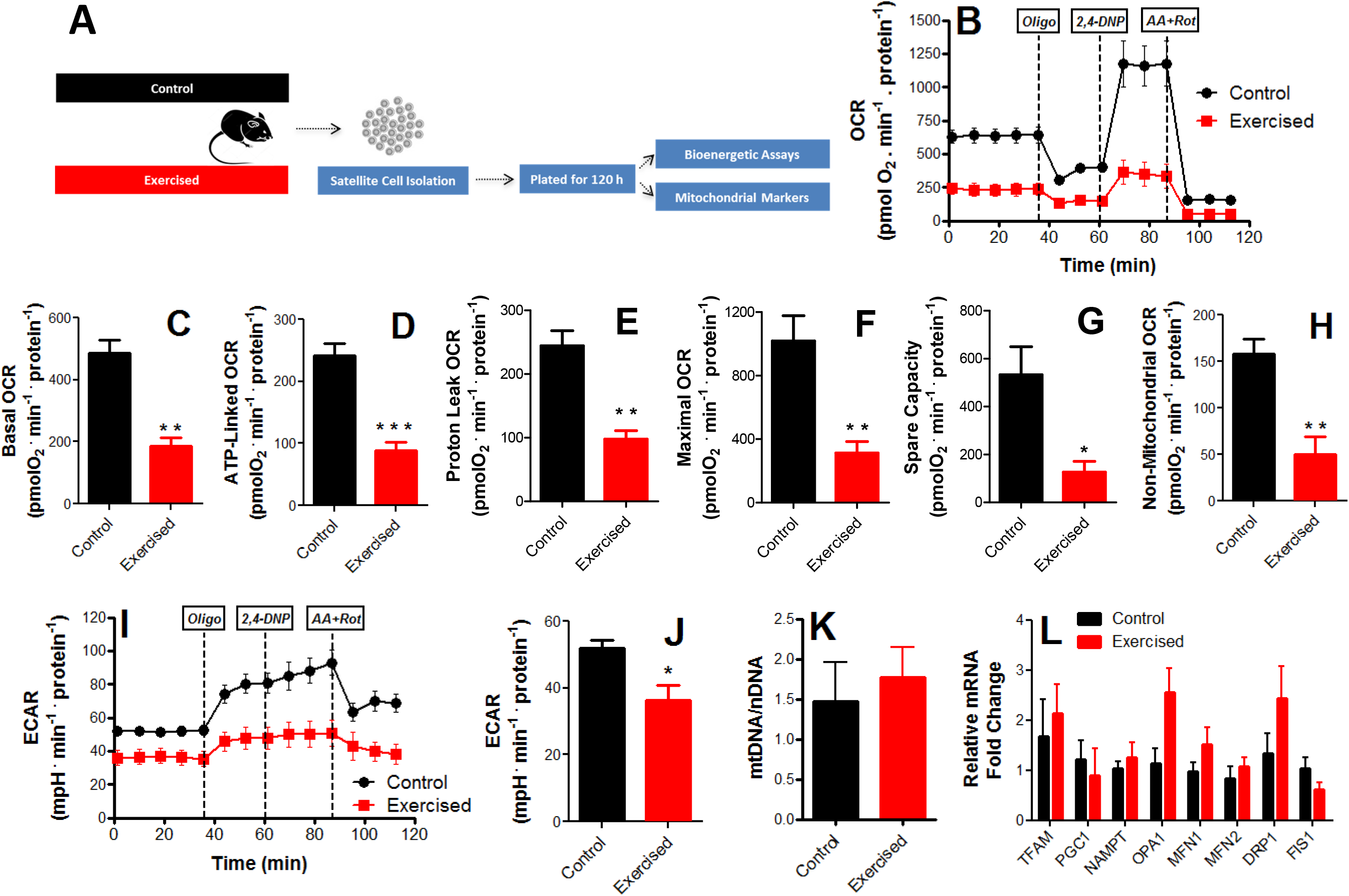
Endurance exercise reduces mitochondrial O_2_ consumption in skeletal muscle stem cells. A: Schematic panel. B: Typical traces of real-time O_2_ consumption rate (OCR) measurements in control and exercised cells determined using the Seahorse XF analyzer. Oligomycin, 2,4-DNP and rotenone plus antimycin A were added where indicated. C: Basal O_2_ consumption rates. D: ATP-production dependent OCR (basal minus oligomycin-insensitive). E: Proton leak (oligomycin-insensitive) OCR; F: Maximal OCR in the presence of 2,4 DNP. G: Spare respiratory capacity (maximal minus basal). H: Non-mitochondrial (antimycin plus rotenone-insensitive) OCR. I: Typical extracellular acidification rate (ECAR) measurement. J: ECAR quantification. K: Mitochondrial DNA/nuclear DNA ratio (mtDNA/nDNA), L: Expression profile of mitochondrial markers. Data represent averages ± SEM and were compared using t tests. Asterisks denote statistically significant differences (*p<0.05, **p<0.01, ***p<0.001) versus control.

A typical trace of real-time O_2_ consumption rate (OCRs) measurements is show in Fig. 5B. Endurance exercise promoted overt changes in mitochondrial bioenergetics, with a substantial suppression of basal, physiological, O_2_ consumption rates (initial part of the trace in Fig. 5B, and quantified in Fig. 5C). This was in part due to lower ATP-linked OCRs, or the amount of inhibition promoted by the addition of ATP synthase inhibitor oligomycin (Fig. 5D), in exercised cells. Additionally, oligomycin-insensitive O_2_ consumption (Fig. 5E), which represents the proton leak of the inner membrane, was decreased by exercise. Exercised cells also presented lower maximal respiratory rates, prompted by the addition of the uncoupler 2,4-dinitrophenol (2,4-DNP, Fig. 5F) and lower spare capacity, or the difference between basal and maximal respiration (Fig. 5G). However, spare capacity was still present, indicating that basal respiratory levels were not limited by overall respiratory capacity. Finally, exercised cells presented less non-mitochondrial respiration, measured in the presence of antimycin plus rotenone (Fig. 5H), although these OCRs were low compared to overall OCRs. Non-mitochondrial OCRs were subtracted from all other OCRs for quantifications.

Inhibition of mitochondrial respiration is often accompanied by a shift toward glucose fermentation to lactate. Indeed, many cells, when more quiescent, are more fermentative (Coller, 2019). However, extracellular acidification rates (ECAR, Fig 5I-J) in exercised cells were significantly decreased in exercised cells, suggesting that they indeed have lower ATP production, and not a shift toward acid-generating lactate formation. Lower ATP demands in the exercised cells are compatible with the expression profile observed previously, suggestive of a higher return to quiescence induced by endurance exercise.

Despite the changes in respiratory capacity, exercise did not alter mitochondrial DNA levels relative to nuclear DNA (Fig. 5K). This indicates that the changes in respiration are not accompanied by changes in mitochondrial mass. Indeed, the expression both of proteins involved in mitochondrial biogenesis and morphology was not significantly altered by exercise (Fig. 5L). Overall, our results show that despite a lack of modulation of mitochondrial mass, endurance exercise induces a repression of mitochondrial respiration in satellite cells, as well as lower energy demands.

Mitochondrial OCRs were found to be decreased by endurance exercise, an effect associated with a more quiescent expression profile. However, oxidative metabolism is important for stem cell proliferation and activation (Coller, 2019, Khacho and Slack, 2017). Consequently, we evaluated satellite cell metabolism in control or exercised animals after activation induced by injury (Fig. 6A). Interestingly, while control OCR values were not significantly altered by injury (compare Figs. 5 and 6B-H), under injured conditions exercised satellite cell OCRs were higher, and equal to that observed in non-exercised cells, under all respiratory states (Figs. 6B-H). ECAR values were also equal between control and exercised activated satellite cells (Fig. 6I-J). This demonstrates that the quiescent metabolic state of exercised cells under non-activated conditions can promptly respond to activating stimuli, with enhanced metabolic activity, as reflected by enhanced mitochondrial ATP production and respiratory capacity. Interestingly, although metabolic activity was equal, the activated exercised cells showed higher colony formation (Fig. 6K), demonstrating that their activated state is more efficient in energy use toward proliferation.

**Fig 6.**
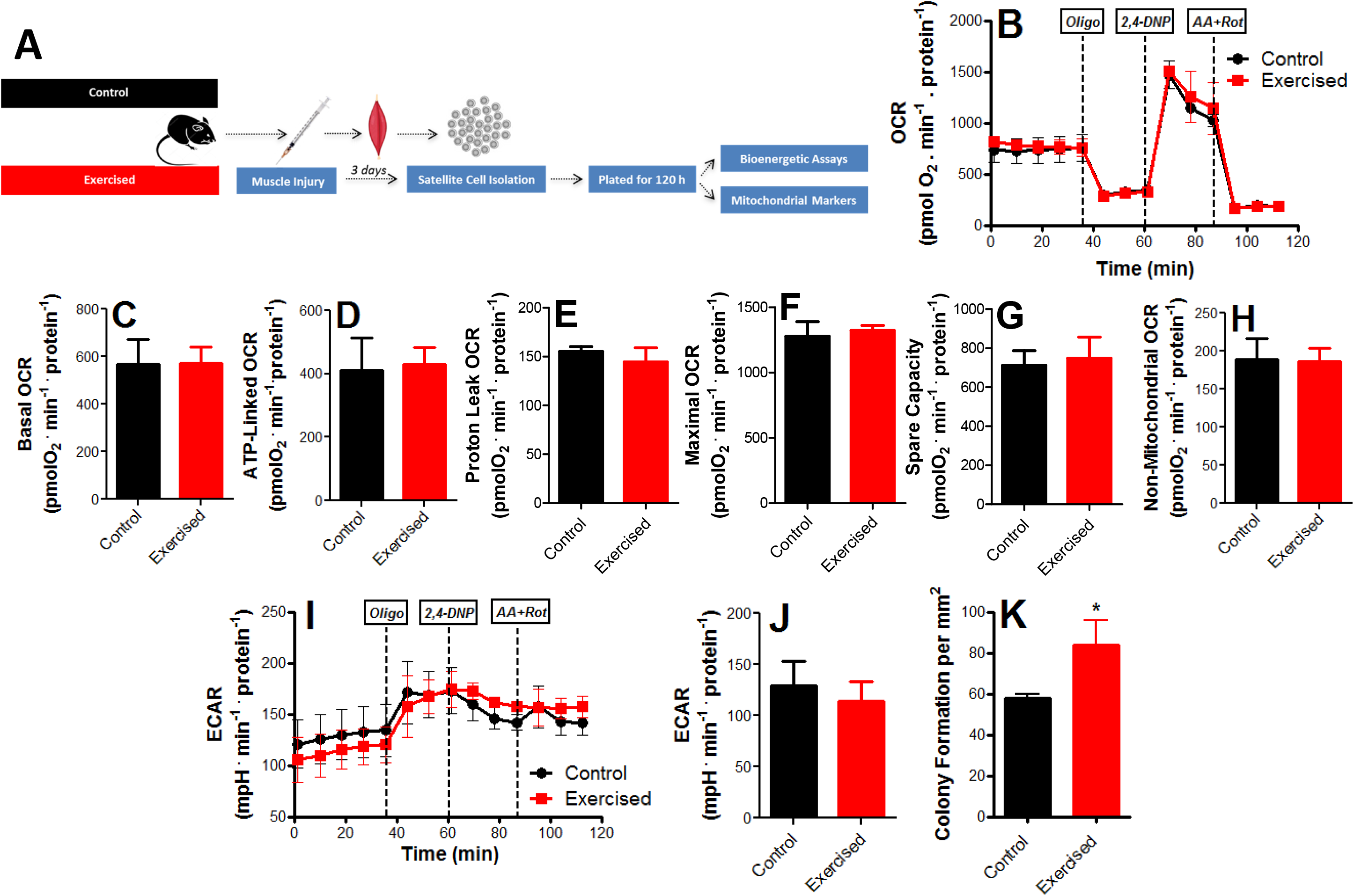
Mitochondrial O_2_ consumption is unaffected in muscle stem cells from injured animals. A: Schematic panel. B: Typical trace. Oligomycin, 2,4-DNP and rotenone plus antimycin A were added where indicated. C: Basal O_2_ consumption rates. D: ATP production-dependent OCR. E: Proton leak OCR; F: Maximal OCR. G: Spare respiratory capacity. H: Non-mitochondrial OCR. I: Typical ECAR trace. J: ECAR quantification. K: Myogenic colony formation. Data represent averages ± SEM and were compared using t tests. Asterisks denote statistically significant differences (*p<0.05) versus control.

### Inhibition of mitochondrial O_2_ consumption increases satellite cell self-renewal and quiescence, while decreasing inflammation upon engraftment

Since endurance exercise promotes mitochondrial respiratory inhibition associated with enhanced return to quiescence and self-renewal markers, we verified if the respiratory inhibition observed was a cause of the changes in expression profile. To do so, we isolated satellite cells from control animals (Fig. 7A), which have high OCRs, and titrated low concentrations of the specific mitochondrial respiratory inhibitor antimycin A, to achieve a respiratory inhibition similar to that observed in exercised animals (Figs. 7B and 7C). Our titrations showed that 70 ng/mL antimycin A promoted respiratory inhibition similar to that observed in exercised cells. Interestingly, when we promoted this respiratory inhibition, the expression profile of non-exercised cells became very similar to that observed in exercised cells, with an increase in stemness, self-renewal and return to quiescence markers (Fig. 7D). This indicates that respiratory suppression is sufficient to promote a phenotype more similar to that of endurance-exercised animals.

**Fig 7.**
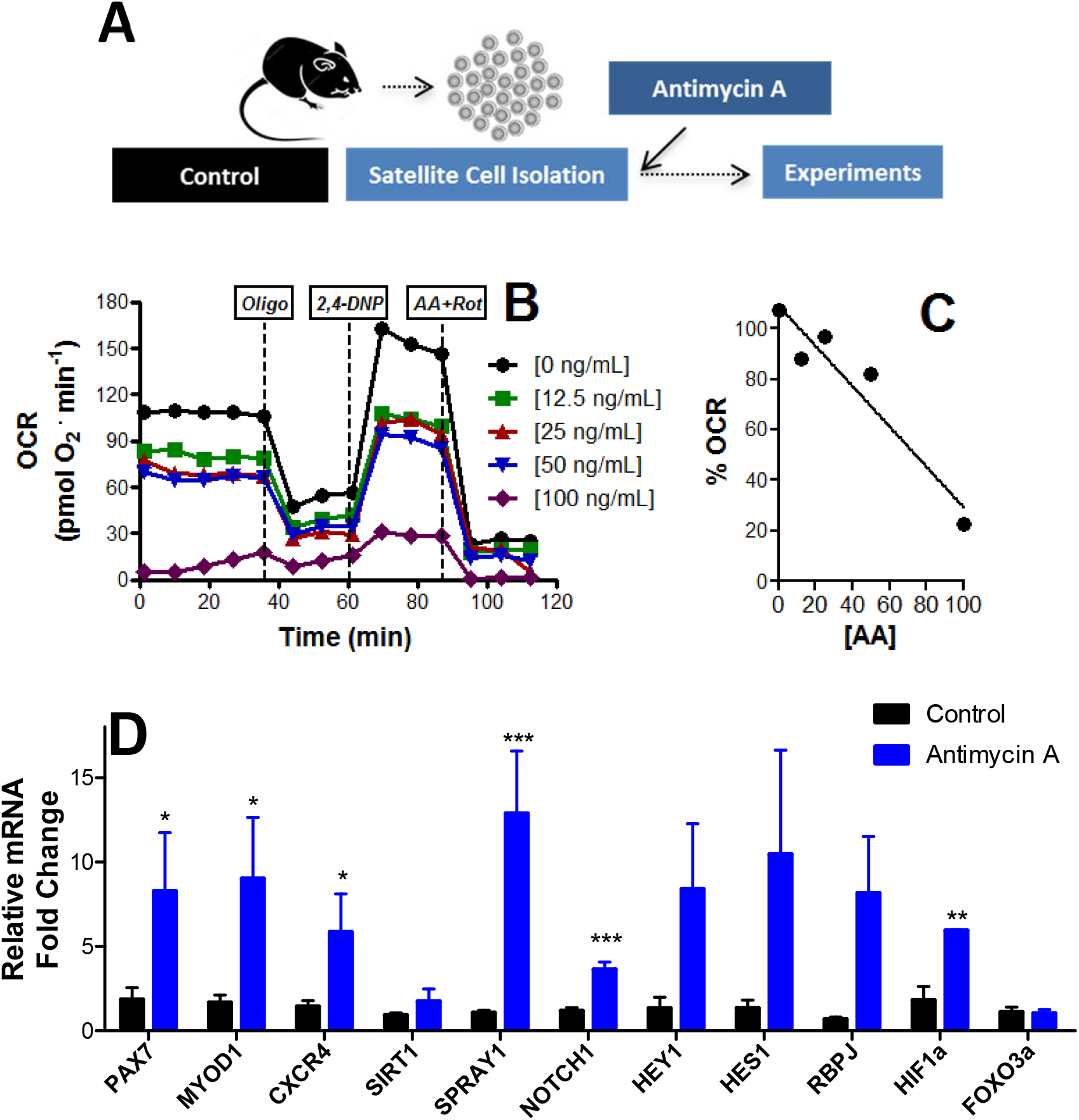
Inhibition of mitochondrial O_2_ consumption promotes satellite cell self-renewal. A: Schematic panel. B: typical OCR trace in the presence of different concentrations of antimycin A. B: Respiratory inhibition versus antimycin A concentrations. C: Satellite cell expression profile after 8 h in the presence of 70 ng/mL antimycin A. Data represent averages ± SEM and were compared using t tests. Asterisks denote statistically significant differences (*p<0.05, **p<0.01, ***p<0.001) versus control.

Next, we examined if exercised or respiratory-inhibited satellite cells can lead to changes in responses when transplanted. To do so, isolated cells from control animals, treated or not with 70 ng/mL antimycin A, or cells from exercised animals were transplanted into control animals which had been previously injured to induce engraftment (Fig. 8A). Illustrative images of tibialis anterior muscles transplanted with control, exercised and antimycin-treated satellite cells are shown in Fig. 8B. While no overt changes in engraftment were noted between treatment groups, as verified by a lack of changes in centralized nuclei (Fig. 8C), both exercised and antimycin-treated cells lead to muscles with significantly lower inflammation (Fig. 8D). This anti-inflammatory effect is most probably related to the enhanced stemness of exercised and respiratory-inhibited satellite cells.

**Fig 8.**
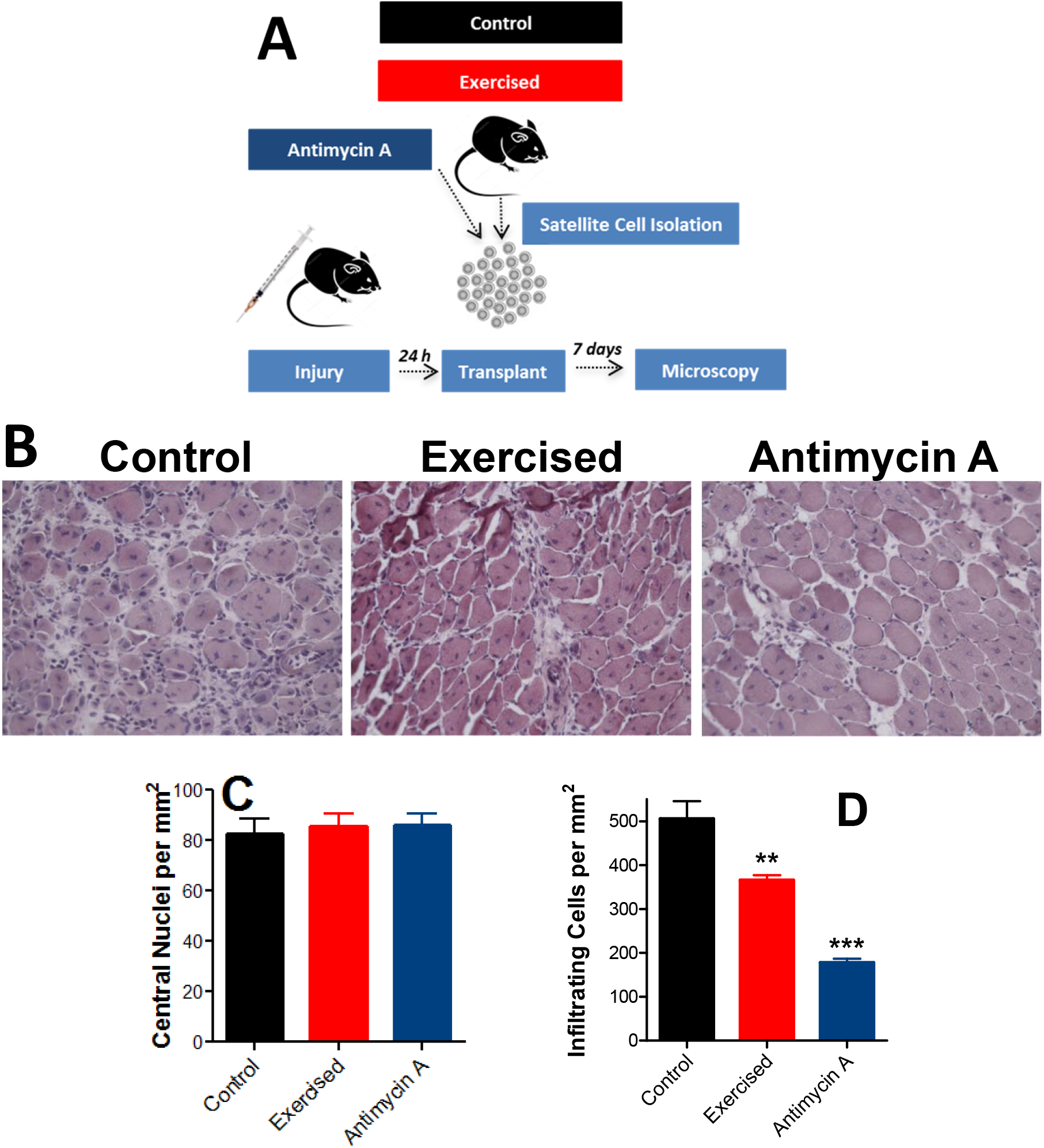
Transplant of exercised or respiratory-inhibited satellite cells prevents inflammation. A: Schematic panel. B: Representative images of transplanted fiber cross-sections. C: Centrally nucleated (newly formed) myofiber quantification. D: Inflammatory cell quantification. The data represent averages ± SEM and were compared using t-tests. Asterisks denote statistically significant differences **p<0.01, ***p<0.001) versus control animals.

## Discussion

Endurance exercise is an often-practiced form of exercise that significantly increases skeletal muscle energy and oxygen demands, thus changing metabolic regulation in the whole body (Hawley et al., 2014; Wilkinon et al., 2008). Indeed, it is a well-established intervention to preserve muscle function and prevent changes in energy metabolism, such as insulin resistance, that occur with aging (Lanza et al., 2008). Despite its widespread use, the cellular and molecular mechanisms involved in metabolic changes promoted by endurance exercise still remain to be fully understood.

Using a mouse model of endurance exercise involving 5 weeks of daily treadmill runs, we found, predictably, that endurance exercise increases physical performance (Fig. 1). This occurred in parallel with changes in oxygen consumption at rest and when exercised, and changes in exhaustion and recovery glucose levels, confirming exercise induces a shift in whole body metabolism. Indeed, exercised animals were leaner, as indicated by lower fat areas and body mass (Fig. 2). Perhaps counter-intuitively, endurance exercise did not change muscle areas or fiber cross sectional areas. These results are in agreement with early work on metabolic effects of exercise, which show that endurance does not promote muscular hypertrophy, but leads to enhanced respiratory and lipid oxidation capacity, thus sparing the body from carbohydrate depletion (Molé et al, 1971; Holloszy and Booth, 1976). These early results indicating glucose conservation is a mediator of enhanced stamina in endurance exercise have been confirmed using modern flux analysis techniques (Overmyer et al., 2015). Recently, peroxisome proliferator-activated receptor delta (PPARδ) was found to be a key mediator in the metabolic shift promoted by endurance exercise, increasing palmitate oxidation without an overall change in mitochondrial biogenesis in myocytes (Fan et al. 2017).

Although no muscular hypertrophy was observed in exercised animals, prior studies have demonstrated that aerobic exercise changes satellite cell numbers and function, thus enhancing muscle regenerative capacity and preserving muscle mass during aging (Kurosaka et al. 2009; Shefer et al. 2010; Cisterna et al. 2016). Indeed, we found that injured muscles in endurance-exercised animals had improved regenerative capacity, as indicated by a higher number of centrally-located nuclei, lower counts of inflammatory cells and less fibrosis (Fig. 3). This improved regenerative capacity may be seminal toward skeletal muscle maintenance in aging, which is enhanced by endurance training (Shefer et al. 2010; Cisterna et al. 2016).

To uncover changes in satellite cells associated with the increased regenerative capacity observed in endurance exercise, we isolated these cells using the preplating method (Gharaibeh et al. 2008). We found that endurance exercise promoted an increase in return to quiescence and self-renewal markers, as well as more undifferentiated morphology, but no changes in colony-forming capacity (Fig. 4). Self-renewal and return to quiescence are pivotal toward maintenance of the satellite cell pool (reviewed in Feige et al., 2018; Forcina et al., 2019), and may explain why endurance exercise promotes more favorable repair after injury.

Interestingly, changes in energy metabolism, and particularly mitochondrial function, have been shown to be central regulatory points in the cell cycle. In many cell types, less differentiation is accompanied by low aerobic metabolism and high glycolytic rates (Varum et al., 2011; reviewed in Xu et al., 2013). However, stem cell activation and differentiation is by no means a homogeneous process, and some final cell fates involve decrease of mitochondrial respiratory activity upon commitment to differentiation (Forni et al., 2016). Furthermore, many studies suggesting a shift from fermentation to oxidative metabolism during differentiation are based on gene expression patterns, while energy metabolism is mostly regulated post-transcription, so gene expression patterns may differ substantially from functional assays (de Carvalho et al., 2017).

As a result, we measured metabolic fluxes in satellite cells isolated from control or endurance exercised mice. Our results show a clear repression of oxidative metabolism in exercised cells, which present lower mitochondrial oxygen consumption rates under all respiratory states (Fig. 5), despite a lack of changes in markers of mitochondrial mass, biogenesis and dynamics. The lower oxygen consumption rates are not attributable to higher rates of fermentation to lactate, as indicated by lower extracellular acidification rates, and are instead related to lower ATP demands. Indeed, respiratory rates in satellite cells isolated from injured muscles were equal in control and exercised cells (Fig. 6), showing that exercise did not ablate the ability to respond to injury. Overall, these results suggest that exercise preserves a more quiescent state in un-activated satellite cells, without compromising their activation response. Higher self-renewal and quiescent capacity may be important for responses toward injury, as indicated by the fact that more newly formed fibers were observed in injured exercised animals (Fig. 3), and colony formation was more pronounced in satellite cells isolated from exercised animals after injury (Fig. 6).

Previous studies have analyzed the role of oxidative mitochondrial metabolism in satellite cells. Zhang et al., (2016) found that aged satellite cells had lower oxidative phosphorylation. However, this was not associated with enhanced quiescence, but instead senescence, as is perhaps expected in aging. Indeed, the loss of satellite cell function with aging could be reversed by recovering NAD levels, in a Sirt1-dependent manner. Consistently, Sirt1 ablation leads to premature differentiation (Ryall et al., 2015), a result compatible with our finding that endurance exercise enhances Sirt1 expression, as well as prevents differentiation and promotes self-renewal. Interestingly, caloric restriction, an intervention that leads to enhanced Sirt1 expression in satellite cells (Cerletti et al., 2012), does not promote other alterations seen in our exercised satellite cells (depressed respiratory rates in the absence of increased mitochondrial DNA), but instead seems to increase oxidative metabolism and enhance mitochondrial biogenesis. This suggests satellite cell remodeling by exercise and dietary restriction differ mechanistically, although both interventions help preserve muscle mass during aging.

Since we observed that exercised satellite cells had lower respiration and higher self-renewal, we questioned if the return to quiescence characteristics induced by exercise were secondary to the lower metabolic rates. Indeed, suppression of oxidative phosphorylation often promotes self-renewal (reviewed in Khacho and Slack, 2017). To achieve this, we promoted respiratory inhibition in control satellite cells to similar levels of those observed in exercised cells by adding low quantities of the classic respiratory inhibitor antimycin A. Remarkably, respiratory suppression alone was sufficient to induce an expression panel in satellite cells similar to that promoted by exercise, with enhanced quiescence and self-renewal markers (Fig. 7).

We tested next if satellite cells with modified by exercise or respiratory modulation could exhibit changes in engraftment after transplant such as those seen in calorically-restricted satellite cells (Cerletti et al., 2012). Interestingly, neither exercised nor respiratory-inhibited satellite cells increased centrally-localized nuclei when transplanted into control animals after injury (Fig. 8). This suggests that the regenerative potential of exercised cells observed *in vivo* (Fig. 3) may depend on characteristics not only of the cells but also their particular niche, and thus are not reproduced upon transplantation. However, consistently with their enhanced quiescence markers, both exercised and respiratory-inhibited satellite cell transplant significantly decreased inflammatory response. This demonstrates that at least part of the beneficial effects of exercise on muscle satellite cell responses can be mimicked by partially suppressing mitochondrial oxidative phosphorylation.

Overall, we demonstrate here that endurance exercise promotes changes in satellite cell function, stemness, self-renewal and differentiation. The changes are associated with repression of mitochondrial oxygen consumption, and artificial suppression of respiration in satellite cells mirrors the characteristics of exercise. Our study provides insights into mechanisms governing myogenesis promoted by exercise that will hopefully contribute toward better therapeutic interventions promoting regulated myogenesis and prevention of sarcopenia.

## Acknowledgments

The authors thank Camille Caldeira for exceptional lab management, Prof. José Cesar Rosa Neto for use of laboratory installations, Dr. Matheus Mori for help with mDNA measurements and Dr. Angela Castoldi for help with the calorimeter. This research was supported by the *Fundação de Amparo à Pesquisa do Estado de São Paulo* (FAPESP), grant number 2016/18633-8, *Conselho Nacional de Pesquisa e Desenvolvimento* (CNPq) grant number 440436/2014, *Coordenação de Aperfeiçoamento de Pessoal de Nível Superior* (CAPES) finance code 001, and the *Centro de Pesquisa, Inovação e Difusão de Processos Redox em Biomedicina* - CEPID Redoxoma, grant 2013/07937-8.

## Author Contributions

Conceptualization: Phablo Abreu and Alicia Kowaltowski

Data curation: Phablo Abreu and Alicia Kowaltowski

Formal analysis: Phablo Abreu

Funding acquisition: Phablo Abreu and Alicia Kowaltowski

Methodology: Phablo Abreu and Alicia Kowaltowski

Project administration: Phablo Abreu and Alicia Kowaltowski

Supervision: Alicia Kowaltowski

Writing – original draft: Phablo Abreu and Alicia Kowaltowski

Writing – review & editing: Phablo Abreu and Alicia Kowaltowski

---

**Supplementary Table 1:**
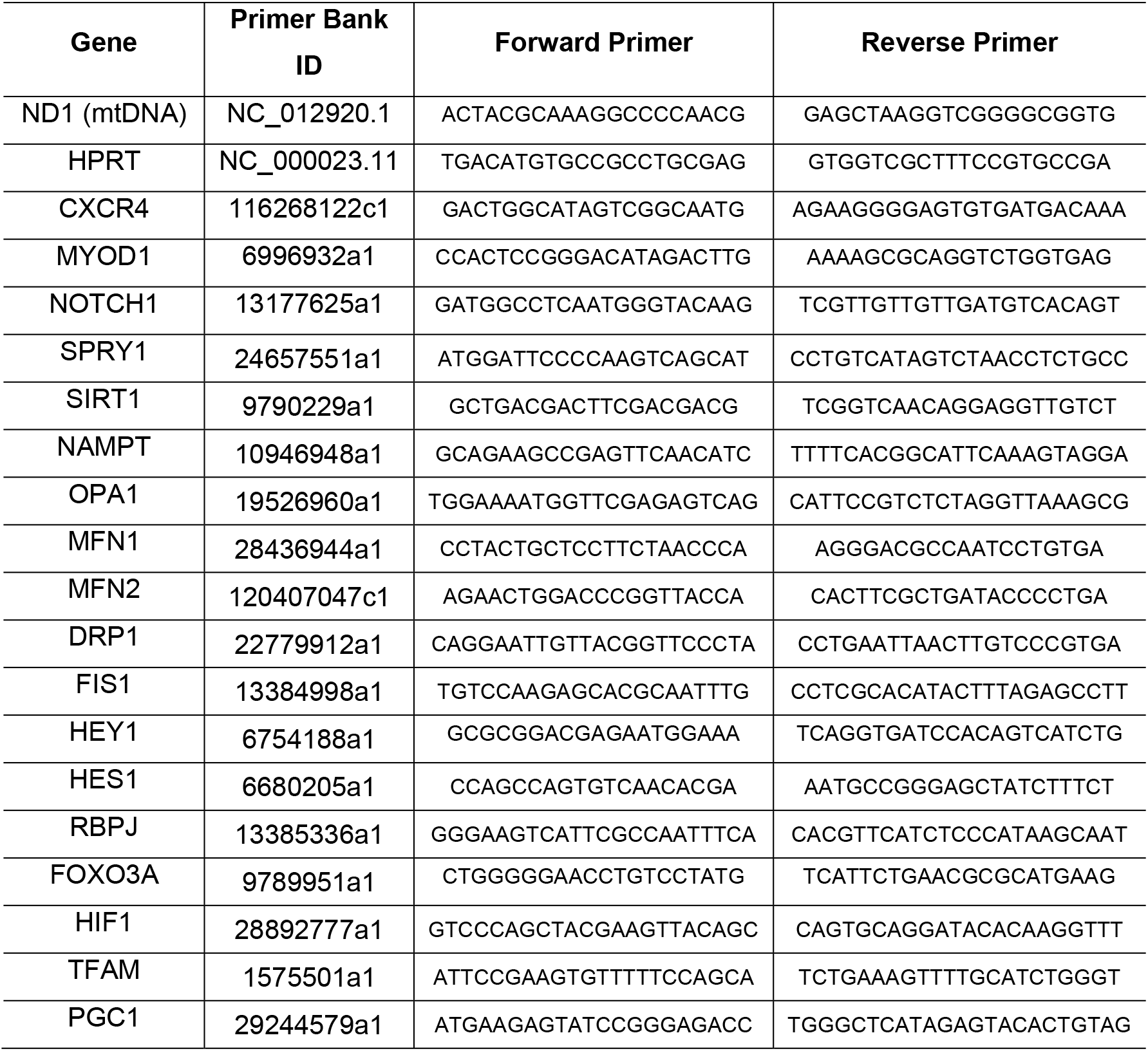
Genes and primers

